# mRNA-1273.251 and mRNA-1283.251 vaccines expressing SARS-CoV-2 variant LP.8.1 antigens broadly neutralize contemporary JN.1-lineage viruses

**DOI:** 10.1101/2025.09.16.676634

**Authors:** Arshan Nasir, Diana Wing Lee, Ziqiu Wang, Clinton Ogega, Nathen Bopp, Phylicia Wisti, Dan Makrinos, Hardik Jani, Amy Gorrie, Michael F Bender, Adrian R Laciak, Arvind Sharma, Xiaozhen Hu, Tessa Speidel, Fadi Ghantous, Daniela Montes Berrueta, Yixuan Jacob Hou, Kath Hardcastle, Sayda Elbashir, Cathal Harmon, Nicholas J Amato, Kai Wu, William R Scheif, Kiera Hohn, Andrea Carfi, Yadunanda Budigi, Rahnuma Wahid, Darin K Edwards, Alec W Freyn

## Abstract

The continued evolution of the SARS-CoV-2 Omicron JN.1 lineage has led to the emergence of antigenically distinct subvariants including KP.2, KP.3, XEC, and LP.8.1, which became the dominant strains in the Americas and Europe by mid-2025. LP.8.1 was designated a Variant Under Monitoring by the WHO in January 2025 due to its potential to displace prior circulating variants. Informed by early growth modeling and antigenic analysis, we selected LP.8.1 as a candidate strain for the 2025-2026 vaccine season. Here, we describe the development of updated LP.8.1-matched mRNA vaccine compositions encoding either the full-length spike protein for mRNA-1273 (monovalent) or the membrane-anchored receptor-binding and N-terminal domains for the mRNA-1283 vaccine. Initial *in vitro* characterization, including structural analysis, demonstrated robust antigen expression and intact antigenic features. Immunogenicity of both vaccines were evaluated in murine models following immunization as either a primary series in naïve animals or as a booster dose. LP.8.1-matched vaccines elicited strong neutralizing antibody responses against the homologous LP.8.1 strain and more recently emerging JN.1-lineage subvariants, including XFG and NB.1.8.1. Notably, the mRNA-1283 vaccine expressing LP.8.1 induced higher mean neutralization titers than the mRNA-1273 version across multiple variants. These data demonstrate the immunogenicity and breadth of both LP.8.1-based mRNA-1273 and mRNA-1283 vaccines in the context of ongoing JN.1 lineage evolution and support the selection of LP.8.1 as the updated vaccine antigen for the 2025-2026 season.

## Introduction

The Severe acute respiratory syndrome coronavirus 2 (SARS-CoV-2) virus continues to evolve largely through stepwise mutations in the spike (S) gene and recombination between co-circulating lineages. This ongoing diversification, particularly within the Omicron JN.1 lineage, has led to the emergence of subvariants with increased potential for immune escape and transmission. As a result, regular updates to vaccine antigen composition remain critical for optimal protection against circulating strains. Since 2022, variant-adapted mRNA-1273 vaccines have been authorized/approved including the bivalent ancestral and BA.4/BA.5 vaccine (mRNA-1273.222) in 2022–2023, and monovalent vaccines targeting XBB.1.5 (mRNA-1273.815) in 2023-24, KP.2 (mRNA-1273.712) and JN.1 (mRNA-1273.167) in 2024–2025, and LP.8.1 (mRNA-1273.251) for the 2025-26 season. Additionally, mRNA-1283.251, the next-generation vaccine expressing membrane-bound Spike Receptor Binding Domain (RBD) and N-terminal Domain (NTD) as a single antigen, was approved for use in the US for the 2025-26 season.

LP.8.1 is a descendant of JN.1 virus that was designated a variant under monitoring by the WHO on January 24, 2025, due to its epidemiological relevance and improved fitness over other competing strains at that time.^1^ LP.8.1 emerged in late 2024 but displaced XEC to become the dominant strain in the Americas and Europe by early 2025. In the United States, LP.8.1 accounted for 58% of COVID-19 infections by May 24, 2025.^2^ Although the KP.2 and JN.1-adapted vaccines provided neutralizing responses against LP.8.1, titers were decreased relative to earlier JN.1-lineage strains, indicating ongoing antigenic evolution.^3^ These data prompted the US FDA, WHO, European Medicines Agency, and other global health agencies to recommend LP.8.1 as a suitable vaccine strain for Fall 2025-26 vaccination.^4–6^

Here, we outline the process by which the LP.8.1 strain was selected for preclinical evaluation for the 2025–2026 season. This included antigen characterization of recombinant and cell-surface expressed Spike and RBD/NTD antigens and detailed structural analysis of the LP.8.1 Spike. mRNA vaccine constructs representing mRNA-1273 and mRNA-1283 for LP.8.1 were generated, and immunogenicity was assessed in murine models both in naïve animals and in the presence of pre-existing immunity to SARS-CoV-2, simulating real-world vaccination scenarios. In addition to estimating the binding and neutralizing antibody responses, we also evaluated T cell responses in both primary and booster immunization contexts and assessed the breadth of cellular immunity against multiple prior SARS-CoV-2 Spike variants. Additionally, we report on the neutralization capacity of LP.8.1-based vaccines against currently circulating variants XFG and NB.1.8.1 that are now increasing in frequency, to evaluate the ongoing relevance of LP.8.1 as a vaccine component. These data support the use of LP.8.1 in the evolving SARS-CoV-2 landscape and demonstrate the value of our preclinical, forward-looking approach to strain selection and annual COVID-19 vaccine composition updates.

## Results

### Epidemiologic analysis and structural characterization of the SARS-CoV-2 LP.8.1 variant

In 2024, further diversification within the JN.1 lineage gave rise to next-generation variants such as KP.2, KP.3.1.1, XEC, and LP.8.1, each of which successively became dominant and replaced older strains (Figure 1A). LP.8.1 emerged in late 2024 and quickly displaced XEC to become the dominant strain by early to mid 2025. More recently, newer subvariants of JN.1 (NB.1.8.1, and XFG) have dominated the variant landscape. Importantly, all of these lineages are within the JN.1 family (Figure 1B). The Spike protein of the LP.8.1 variant is characterized by nine amino acid mutations relative to JN.1, including four in the RBD (R346T, V445R, F456L, and Q493E; Figure S1A). Two of the three new mutations in the NTD, S31-(a deletion) and R190S, both introduce new N-linked glycosylation motifs on the LP.8.1 Spike and are predicted to impact the conformation of the RBD. Although our risk assessment model predicted a 2.97 and 1.68 fold-drop for LP.8.1 variant against JN.1 and KP.2 vaccine or infection-derived neutralizing antibodies (nAbs) (Figure S1B), LP.8.1 was prioritized (based on its increasing global growth) for at-risk vaccine preparation as a potential composition for the 2025/2026 season (named mRNA-1273.251).^7^

**Figure 1.**
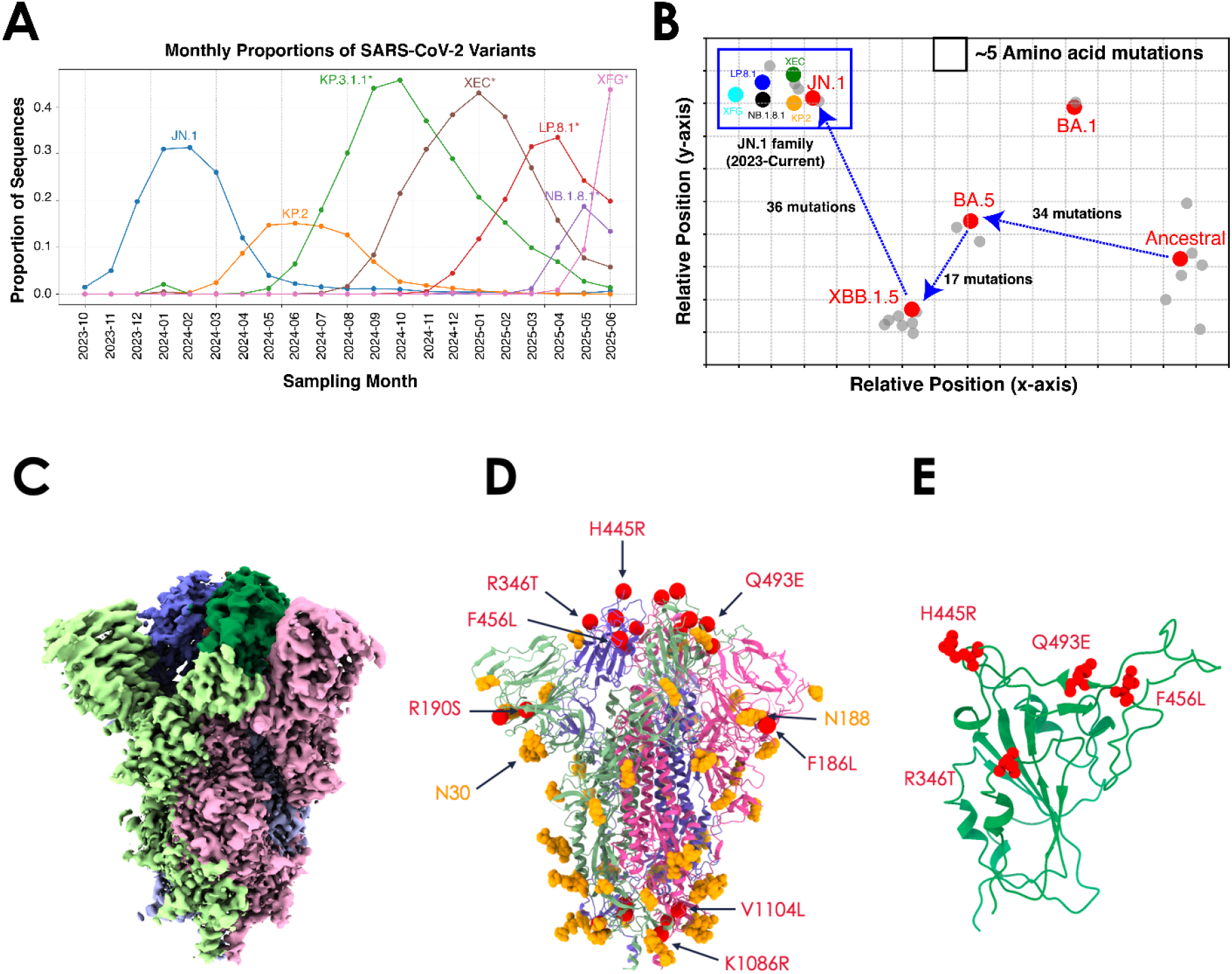
Epidemiologic and structural analysis of SARS-CoV-2 variants. (A) Global dynamics of SARS-CoV-2 variants from November 2023 through June 2025, based on sequences submitted to public databases and analyzed using the nextclade algorithm (ver. 3.15.2). (B) Genetic cartography illustrates the S amino acid distance among SARS-CoV-2 variants. Each grid square corresponds to ∼5 mutations. JN.1 lineage variants are highlighted inside a blue box. (C) Side view of the CryoEM density map of the SARS-CoV-2 LP.8.1 spike ectodomain. Each protomer is colored distinctly (pink, green, and blue), with the RBD of each protomer highlighted in a darker shade. (D) Refined atomic model of the LP.8.1 spike highlighting mutations relative to JN.1 (red or orange spheres). N-linked glycans (orange); sites differing from JN.1 are labeled. (E) Zoomed view of RBD with mutations highlighted.

Structural studies of the LP.8.1 variant spike protein were conducted to characterize the structure and conformation of the new variant spike. A single-particle structure of the unliganded Spike was determined by cryo-electron microscopy at 2.6 angstrom resolution. The structural analysis revealed that the majority of the particles present the RBD subdomains in the “down” conformation, rather than in mixed “up” and “down” conformations as have been seen with previous Spike variants including JN.1 (Figure 1C, S2A, and Table S1).^8–12^ Mutations that differ between the LP.8.1 variant and the prior year JN.1 variant are found on the surface of the spike across multiple domains (Figure 1D and 1E).

More recently, JN.1-lineage subvariants NB.1.8.1 and XFG have emerged and are increasing in frequency in Asia and US/Europe, respectively. NB.1.8.1 is derived from the XDV variant that was a recombinant between two JN.1 strains with seven new spike substitutions including four in the RBD (Figure S1A). XFG is another recombinant of two JN.1 variants with a recombination breakpoint inside the RBD. The XFG RBD, partly derived from LF.7.9 and LP.8.1.2, encodes six RBD mutations relative to the JN.1 parental spike (Figure S1A). Analysis of the mutations relative to LP.8.1 has revealed the surfaces at which these mutations reside and inform the risk of immune escape and conformational consequence of the additional mutations. Our *in silico* risk assessment indicated effective neutralization of both NB.1.8.1 and XFG variants following LP.8.1 vaccination (scores of 1.17 and 2.62, respectively; Figure S1B).^7^

### *In vitro* expression and conformational binding characterization of SARS-CoV-2 mRNA constructs

To better characterize our detection reagents for expression analysis, we determined the X-ray crystal structure of Fab 1205 in complex with the SARS-CoV-2 BA.2.86 RBD at 2.5 angstrom resolution (Table S1).^13^ The monoclonal antibody (mAb) 1205’s epitope overlaps the hACE2–RBD interface (PDB: 8XUZ), providing a structural rationale for the antibody’s broad cross-reactivity (Figure 2A and S2B). This overlap supports the use of 1205 as a reliable binding reagent for comparative expression analyses across variants.

**Figure 2.**
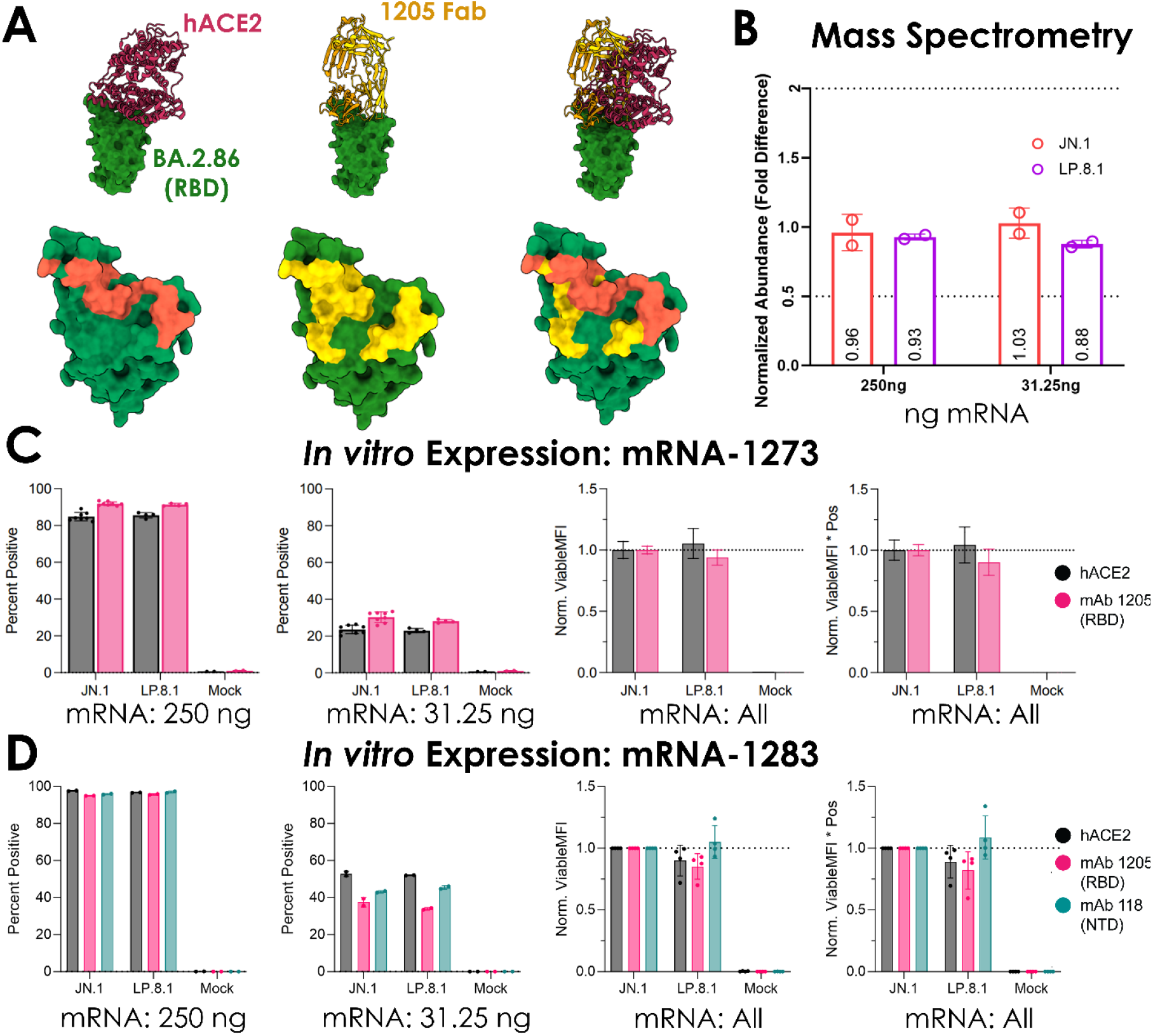
Characterization of mRNA-expressed LP.8.1 antigen. (A) Structures of the SARS-CoV-2 RBD bound to hACE2 (PDB: 8XUZ) and to the broadly neutralizing antibody 1205. The RBD is shown as a green surface; hACE2 and Fab 1205 are shown as red and yellow ribbons, respectively. Top panel: side view. Bottom panel: binding footprints on the RBD. Right panel: superposition highlighting the overlap between the hACE2- and 1205-binding sites. (B) Mass spectrometry showing the relative fold difference in peptide abundance of JN.1 and LP.8.1 constructs after transfection of different mass of mRNA into cells. Biological duplicates are shown as individual dots with the bar representing the mean value plus or minus standard deviation. (C and D) *In vitro* expression of mRNA-1273 and mRNA-1283 (D) versions of JN.1 and LP.8.1 antigens were assessed by flow cytometry after transfection of different masses of mRNA into Expi293F cells. The percentage of positively stained cells are shown after detection with either recombinant human ACE2 (hACE2), 1205, or monoclonal antibody 118 (for mRNA-1283 only). Data for both transfected masses were compiled, and the mean fluorescence intensity (MFI) or MFI multiplied by percent positive cells was normalized to JN.1 to show the detection of LP.8.1 antigen relative to JN.1. Individual values are shown as dots (n=4) with bars representing average values plus or minus standard deviation.

*In vitro* antigen expression in Expi293 cells 24 hours after transfection with codon-optimized SARS-CoV-2 mRNA constructs was assessed by Liquid Chromatography-Mass Spectrometry (LC-MS) for whole cell expression for the mRNA-1273 constructs and flow cytometry for cell-surface expression of both mRNA-1273 and mRNA-1283 constructs. Normalized protein abundance measured by LC-MS showed comparability of JN.1 and LP.8.1 Spike antigen at different mass of mRNA transfected into cells (Figure 2B). Flow cytometry-based analyses using recombinant hACE2 and mAbs targeting RBD (1205) or NTD (118) for detection showed comparable expression between JN.1 and LP.8.1 mRNA-1273 vaccines whether the measure was the percentage of positive cells, the mean fluorescent intensity of viable cells (viable MFI), or the multiplicative product of those two measures (Figure 2C and S3). Analysis of mRNA-1283-based designs using hACE2, 1205 mAb, as well as an additional NTD-specific mAb, also revealed comparable expression levels between JN.1 and LP.8.1 sequences tested regardless of the detection reagent (Figure 2D and S4).

Overall, these data suggest that mRNA-1273 and mRNA-1283 LP.8.1 lineage antigens express comparably to the prior season JN.1 vaccines.

### mRNA-LNP vaccines based on LP.8.1 provide broad protection from JN-1 lineage variants

After confirmation that the appropriate characteristics of the LP.8.1 antigens expressed as full spike antigen (mRNA-1273.251) or as the next-generation membrane-anchored RBD/NTD antigen (mRNA-1283.251), the mRNA were formulated into a lipid nanoparticle for immunogenicity assessment. Studies using the murine model were performed to compare the ability of vaccines with LP.8.1 antigens to elicit binding and functional antibody responses with vaccines from the prior season (JN.1: mRNA-1273.167 and KP.2: mRNA-1273.712) and to assess the breadth of neutralization against JN-1-lineage variants. Mice were dosed in a prime-boost regimen with three weeks interval between doses, and serum was collected two weeks post boost to assess the peak antibody response (Figure 3A). Enzyme-linked immunosorbent assays (ELISAs) to detect binding antibody responses to the original Wuhan-Hu-1 strain were performed (Figure 3B and 3C). Binding antibody titers were comparable (within roughly 2-fold) at both day 21 and 36 timepoints when comparing mRNA-1273 format vaccines (Figure 3B). Immunization with mRNA-1283.251 elicited higher titers (>3-fold) when compared to prior year mRNA-1273-based vaccines after boost (Figure 3C). Neutralization by serum taken at peak titer and two weeks post boost was assessed using a panel of vesicular-stomatitis virus-based pseudoviruses expressing spike glycoproteins matching the variants of interest. Neutralization against LP.8.1 and other currently circulating variants (XFG and NB.1.8.1) was highest after immunization with mRNA-1273.251, matching the LP.8.1 variant, when compared to control JN.1 and KP.2 variant mRNA-1273 vaccines (Figure 3D). When immunized with mRNA-1283.251, the next-generation NTD/RBD design based on LP.8.1, neutralization against all JN.1-lineage pseudoviruses tested was higher than after immunization with JN.1 or KP.2 mRNA-1273 vaccines (Figure 3E).

**Figure 3.**
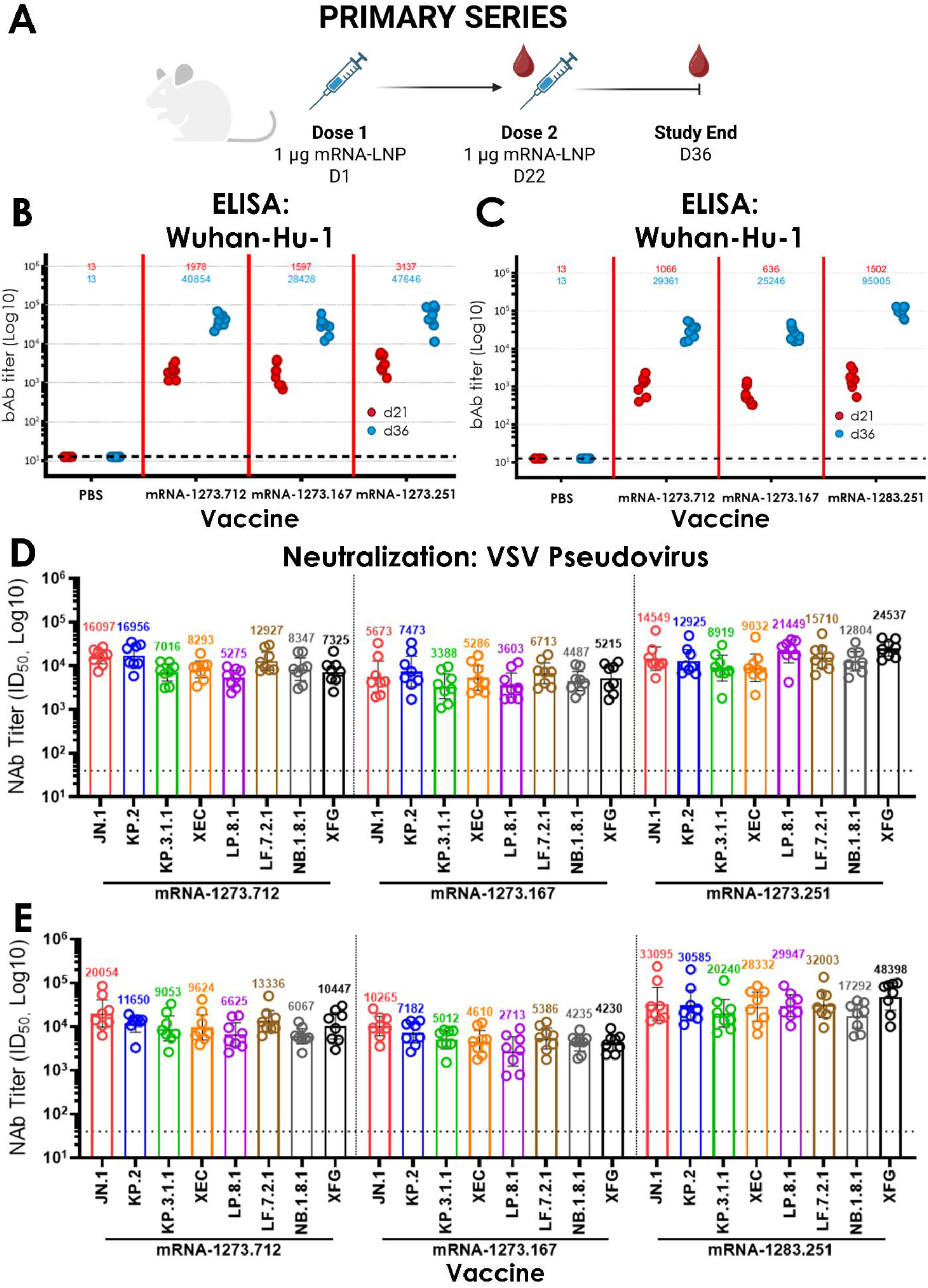
Immunogenicity after Primary Series Immunization with LP.8.1, KP.2, and JN.1-based mRNA-LNP Vaccines. (A) Diagram of the mouse immunogenicity study demonstrating key immunization and sample collection timepoints. (B and C) ELISA binding antibody titers against Wuhan-Hu-1 strain recombinant spike protein comparing animals immunized with mRNA-1273.712 (KP.2) or mRNA-1273.167 (JN.1) with those receiving the (B) full-length spike mRNA-1273.251 (LP.8.1) or (C) NTD/RBD minimal epitope vaccine mRNA-1283.251. GMTs are reported above each group (n=8) and individual animal values represent technical duplicates. (D and E) PsV neutralization titers are reported using peak immune serum two weeks post-boost. Control vaccines were compared to (D) mRNA-1273.251 or (E) mRNA-1283.251. The strain of spike protein expressed on the PsV is described on the x-axis. GMTs are represented by the bar for each group, with the GMT value reported above each bar. Error bars represent the 95% confidence interval and individual data points represent biological duplicates of technical duplicates (n=8/group).

Additional mouse studies were performed to determine the ability of contemporary variant vaccines to stimulate a relevant immune response in the context of diverse, heterologous immunity. Mice were immunized with an initial bivalent dose of mRNA-1273 and mRNA-1273.045 (BA.4/BA.5) then were given a second bivalent immunization with mRNA-1273.815 (XBB.1.5) and mRNA-1273.712 (KP.2) three weeks later. This initial vaccination regimen was designed to elicit immunity in mice that covers the major SARS-CoV-2 lineages that previously circulated and also were vaccine compositions used in licensed vaccines in prior seasons. A booster dose of mRNA-1273.251 (or comparators mRNA-1273.167 or mRNA-1273.712) was administered 43 days later, and serum was collected prior to boost and two weeks post booster dose to assess the peak antibody response (Figure 4A). In a second study, following a similar immunization strategy as above, mRNA-1283.251 was administered as a booster dose (and relevant controls) 66 days after the second primary dose and serum was similarly collected prior to and two weeks post booster dose to assess immunogenicity. ELISAs were used to detect binding antibody responses to the original Wuhan-Hu-1 strain. Antibody titers showed some variability with the pooled pre-boost samples at day 65, but after booster immunization day 79 titers were comparable (with 2-fold) for all mRNA-1273 vaccine variants tested (Figure 4B). Minimal variation was seen with the cohort comparing mRNA-1283.251 to prior year mRNA-1273.167 and mRNA-1273.712 vaccines, with titers within 2-fold change for all groups at both day 88 and day 102 timepoints (Figure 4C). After administration of the heterologous primary series bivalent immunizations, neutralization titers were found to be highest against the more ancestral JN.1-lineage variants (JN.1 and KP.2) and generally decreased with antigenic distance of the pseudovirus variants tested (Figure 4D and 4E). Booster Immunization with mRNA-1273.167 or mRNA-1273.712 raised neutralization titers against all JN.1-lineage pseudoviruses tested, but the pattern of decrease titer with increased antigenic distance from JN.1 and KP.2 held (Figure 4D and 4E). mRNA-1273.251 led to a similar pattern of neutralization as the JN.1 or KP.2 vaccines in the heterologous immunity context (Figure 4D). Vaccination with mRNA-1283.251 showed a more balanced pattern across JN.1-lineage pseudoviruses tested, with titers against all JN.1-lineage pseudoviruses numerically higher than titers generated with all mRNA-1273 variants tested (Figure 4E).

**Figure 4.**
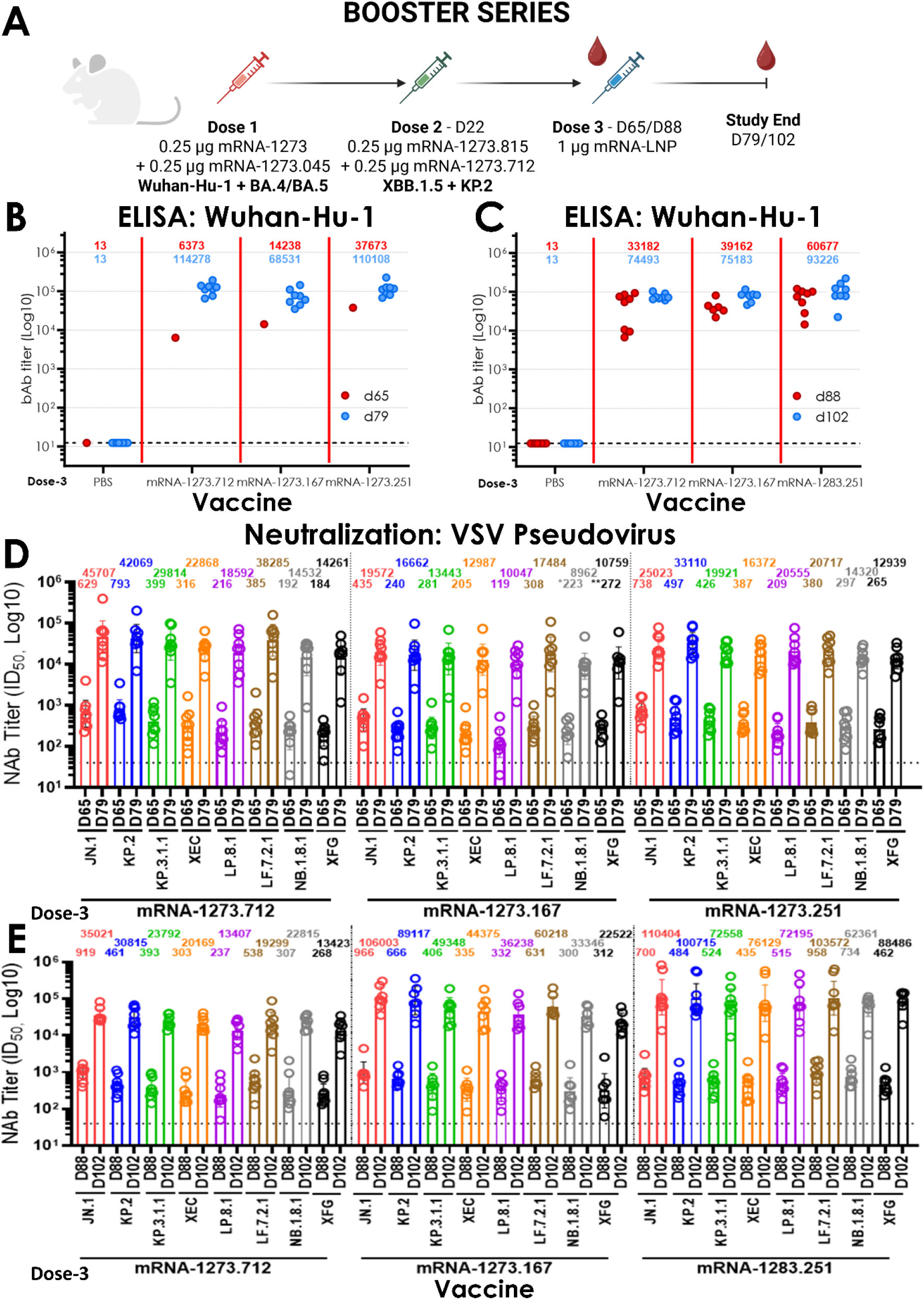
Immunogenicity after Booster Series Regimen with a Heterologous Primary Series. (A) Diagram of the mouse immunogenicity study demonstrating primary series dosing strains as well as key immunization and sample collection timepoints. (B and C) ELISA binding antibody titers against Wuhan-Hu-1 strain recombinant spike protein comparing animals boosted with mRNA-1273.712 (KP.2) or mRNA-1273.167 (JN.1) with those receiving the (B) full-length spike mRNA-1273.251 (LP.8.1) or (C) NTD/RBD minimal epitope vaccine mRNA-1283.251. GMTs are reported above each group (n=8) and individual animal values represent technical duplicates, except pre-boost in B, where a single value is reported representing pooled serum tested in technical quadruplicate. (D and E) PsV neutralization titers are reported using peak immune serum one day pre boost and two weeks post-boost. Control vaccines were compared to (D) mRNA-1273.251 or (E) mRNA-1283.251. The strain of spike protein expressed on the PsV is described on the x-axis. GMTs are represented by the bar for each group, with the GMT value reported above each bar. Error bars represent the 95% confidence interval and individual data points represent biological duplicates of technical duplicates (n=8/group; D - ^*^n=7/group and ^**^n=6/group).

These cumulative data demonstrate the ability of LP.8.1 vaccines to generate neutralization responses that cover currently circulating JN.1-lineage variants with more potency than prior year vaccines.

### Vaccines representing contemporary variants provide broad T cell responses

To determine the potential for contemporary variant vaccines to elicit broad T cell responses, mice were immunized in a primary series regimen with mRNA-1273.251, mRNA-1273.712, or mRNA-1273.167, and spleens were harvested two weeks post second vaccination (Figure 5A). Single-cell splenocyte suspensions were generated and incubated in the presence of overlapping peptides representing the S1 or S2 domain of various SARS-CoV-2 variants from the Wuhan-Hu-1 strain to the contemporary LP.8.1 strain. Peptide stimulation of reactive T cells was measured via intracellular cytokine staining and the interferon gamma (IFN-γ) expressing populations were represented as a function of total CD4 or CD8 T cells measured for each individual animal. In general, CD4 responses were low to both S1- and S2-based peptide pools, with no individual animal eliciting an IFN-γ response higher than 0.5% of total CD4 cells measured (Figure 5B and 5C). Though low, responses were relatively similar across variant-representative peptide pools tested (all averaged between 0 and 0.3% positive CD4 responses). IFN-γ expressing CD8 T cells were robustly stimulated after incubation with S1 peptides, with roughly 2% of CD8 T cells reacting regardless of the variant and vaccine utilized (Figure 5D). Minimal CD8 T cell activity was observed after stimulation with S2-specific peptide pools irrespective of variant or vaccine used (Figure 5E).

**Figure 5.**
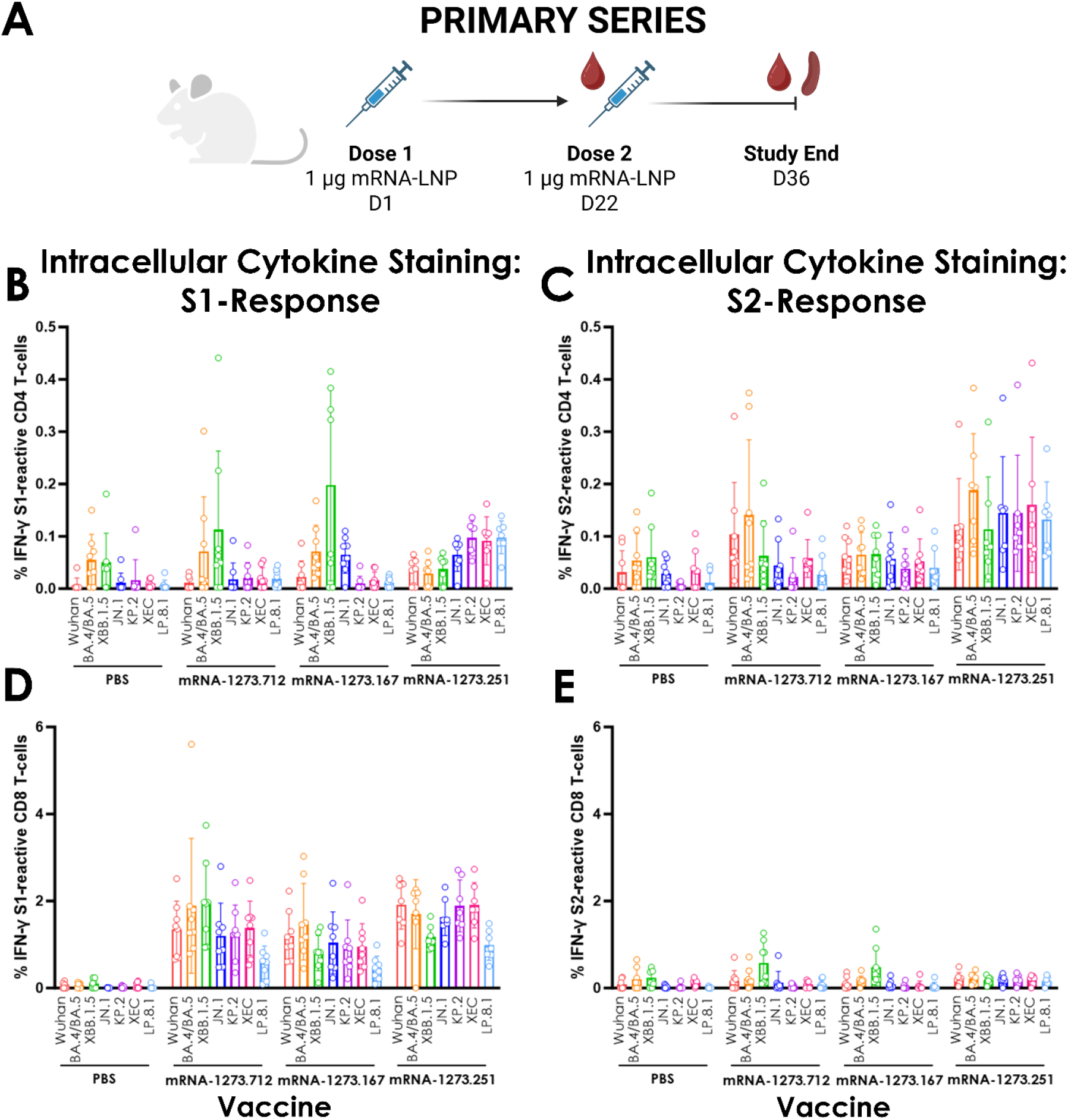
Cellular Immune Response after Primary Series Immunization with LP.8.1, KP.2, and JN.1-based mRNA-LNP Vaccines. (A) Diagram of the mouse immunogenicity study demonstrating key immunization and sample collection timepoints. (B-D) Interferon gamma expression of CD4 (B and C) and CD8 (D and E) T cells after stimulation of splenocyte cell suspensions with overlapping peptide pools representing the S1 (B and D) or S2 (C and E) domains of SARS-CoV-2 spike. Data are shown as percentage of positive cells out of total CD4 or CD8 T cells measured for each individual animal with bars representing the group mean (n=8) and error bars showing standard deviation.

Additionally, mice were immunized with a primary series regimen to generate heterologous immunity that more closely reflects the immune experience of the human population. In this cohort of mice, animals received an initial bivalent dose containing mRNA-1273 (Wuhan-Hu-1) and mRNA-1273.045 (BA.4/BA.5) followed 3 weeks later by a second bivalent dose of mRNA-1273.815 (XBB.1.5) and mRNA-1273.712 (KP.2) (Figure 6A). Both cohorts were boosted with a single dose of a JN.1-lineage variant vaccine (mRNA-1273.167, mRNA-1273.712, or mRNA-1273.251). Two weeks post boost, spleens were harvested and processed into single cell suspensions prior to incubation with overlapping peptide pools representing either the S1 or S2 domain of several SARS-CoV-2 variants. In general, CD4 responses were low to both S1- and S2-based peptide pools, though consistent IFN-γ expression responses were observed across variants, vaccine types, and primary series regimens used (Figure 6B and 6C). Responses were skewed toward Th1 cytokine-expressing cells, rather than Th2 (Figure S5). CD8 T cells responses were strong after incubation with S1 peptides, with around 5% of CD8 T cells showing IFN-γ expression (Figure 6D). S1-reactive CD8 T cells also expressed the activation marker CD107a implying these cells are able to functionally kill infected cells (Figure S6). Approximately, 1% of CD8 T cells reacted to the S2 peptide pools suggesting that the broad initial immunity after primary series enhanced the conserved recall response even after exposure to a novel antigen expressing mRNA-LNP such as mRNA-1273.251 (Figure 6E).

**Figure 6.**
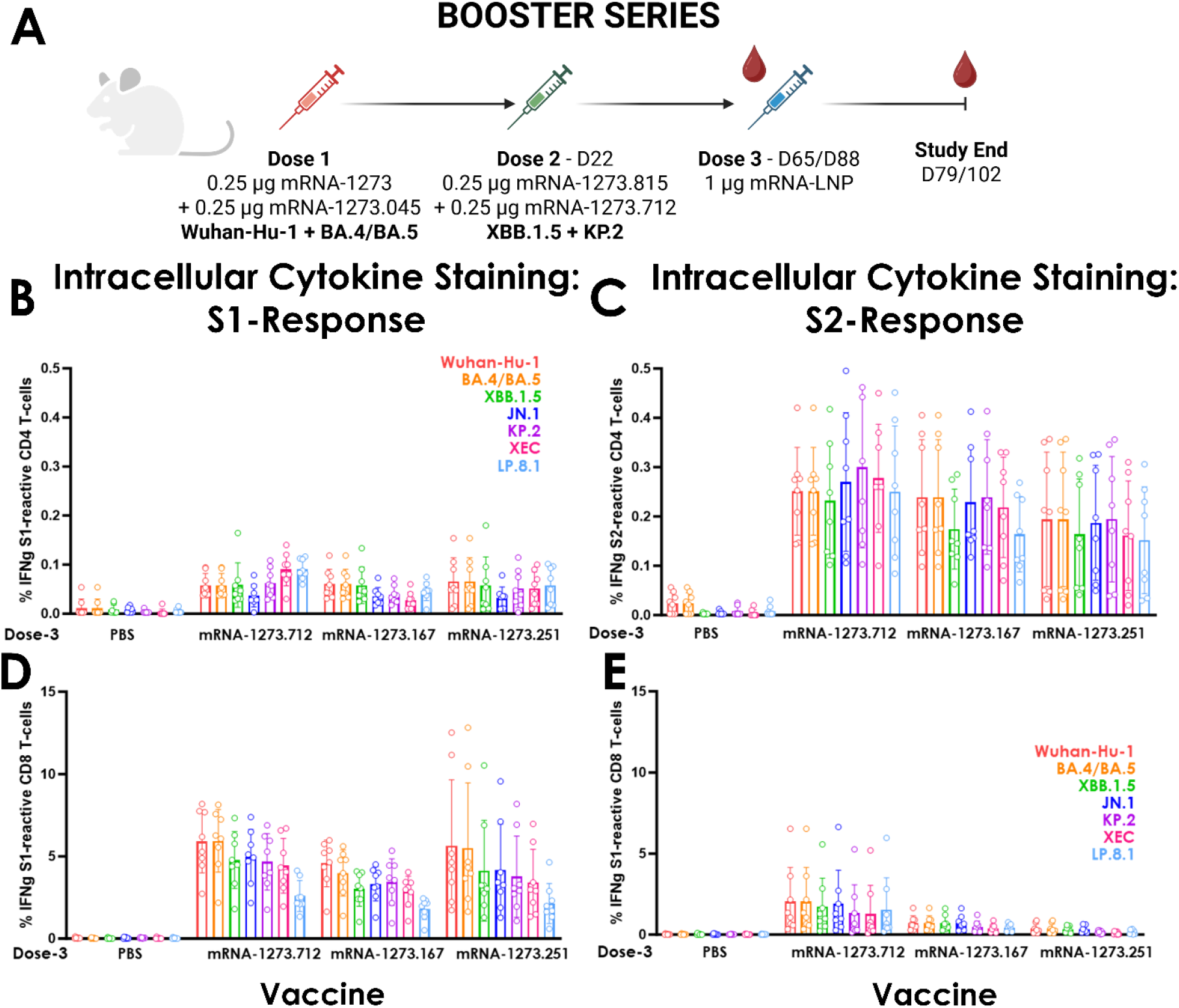
Cellular Immune Response after Booster Series Immunization with a Primary Series of Heterologous Vaccines. (A) Diagram of mouse immunogenicity study demonstrating primary series dosing strains as well as key immunization and sample collection timepoints. (B-E) Interferon gamma expression of CD4 (B and C) and CD8 (D and E) T cells after stimulation of splenocyte cell suspensions with overlapping peptide pools representing the S1 (B and D) or S2 (C and D) domains of SARS-CoV-2 spike. Data are shown as percentage of positive cells out of total CD4 or CD8 T cells measured for each individual animal with bars representing the group mean (n=8) and error bars showing standard deviation.

These data suggest that CD8 T cell responses are stimulated after immunization with mRNA-1273 vaccines and that responses can be observed across variants representing SARS-CoV-2 diversity.

## Discussion

To evaluate the suitability of LP.8.1 for the 2025/2026 vaccine update, a comprehensive preclinical assessment was conducted, including routine variant monitoring, *in vitro* expression analysis of variants of interest, and assessment of immunogenicity in small animal models including functional antibody and T cell responses. Specifically, we analyzed variant growth trends and antigenic profiles against published models of risk assessment to inform early identification of potential strains of interest that showed early signs of strain replacement and immune evasion against approved vaccines.^7^ This exercise led to the selection, in December 2024, of LP.8.1 as a candidate strain for vaccine development for the next season.

Initial *in vitro* characterization was performed with vaccine candidates to ensure the antigen, designed as either full spike (mRNA-1273) or RBD/NTD (mRNA-1283), was effectively processed and expressed on the cell surface. Lead candidates were then moved into murine immunogenicity studies to assess the ability of vaccines to elicit appropriate humoral and cellular immune responses. LP.8.1 vaccines were compared to prior season JN.1 and KP.2-based mRNA-1273 vaccines and elicited robust neutralizing responses covering all JN.1-lineage pseudoviruses tested. Additionally, two contemporary circulating variants emerged after strain selection (XFG and NB.1.8.1), and we found that the LP.8.1 vaccine-elicited responses could neutralize these variants with higher potency than either the JN.1 or KP.2-based vaccines. These findings support the selection of LP.8.1 as the vaccine strain for the 2025/2026 season, offering coverage of the dominant circulating variant and broader protection against emerging variants.

Cellular responses were also investigated after immunization with JN.1-, KP.2-, or LP.8.1-based mRNA-1273 vaccines and found to be broad and potent. The key T cell epitopes known to be present on the spike protein explain the breadth observed for these peptide pools, but responses to the S1 domain also showed responses in both primary and booster studies. Conserved CD8 T cell stimulating peptides in the S1 domain must be present in all strains to provide this broad response in mice, which may be extrapolated to the human cellular response.^14^ As variants change through antigenic drift and lineage shifts, T cell responses are maintained to conserved regions in the Spike protein potentially providing some amount of protection in the absence of a strain-matched neutralizing antibody response.

The totality of data presented here, in combination with data generated by public health groups,^1,15,16^ governments,^2,17^ academic researchers,^18^ and other industry groups supported the use of LP.8.1 as a vaccine antigen for the 2025/2026 season.^4–6^

## Materials and Methods

### Sequence Analysis

SARS-CoV-2 genetic sequences from the Global Initiative on Sharing All Influenza Data (GISAID) database were downloaded and analyzed via the Nextclade algorithm (current version 3.15.2) for viral genome alignment, quality control checks, mutation calling, and clade assignment. S mutations were called relative to the ancestral Wuhan-Hu-1 strain (GenBank Accession ID: MN908947) using in-house Python (version 3.12.10) scripts. Filtered S variants were categorized using risk scoring algorithm and variants that indicated either increased immune-evasion or strain-replacement were prioritized for pseudovirus production and preclinical assessment.^7^

### Structural characterization of LP.8.1 spike via single particle cryo-electron microscopy

Cryo-electron microscopy (cryoEM) grids were prepared by applying 4 μL of purified LP.8.1 spike protein at a concentration of 0.5 mg/mL onto glow-discharged Quantifoil R1.2/1.3 300-mesh copper grids. Grids were blotted and plunge-frozen into liquid ethane using a Vitrobot Mark IV (Thermo Fisher Scientific) set to 4 °C and 100% relative humidity. Data were collected using a Titan Krios G3i transmission electron microscope (Thermo Fisher Scientific) operating at 300 kV and equipped with a Gatan K3 direct electron detector and a BioQuantum energy filter (Gatan) operated in zero-loss mode. Automated data acquisition was performed using EPU software at a nominal magnification of 105,000×, corresponding to a calibrated pixel size of 0.822 Å. A total electron dose of 51.5 e−/Å^2^ was fractionated over 50 frames per movie. The nominal defocus range was set from –0.6 μm to –2.4 μm.

All movie frames were imported and processed in CryoSPARC v4.7.1. Motion correction and CTF estimation were applied to the raw movies prior to particle picking. Particles were automatically picked using Topaz, followed by iterative rounds of 2D classification to discard junk particles and select high-quality projections. 3D classification was employed to assess conformational heterogeneity and remove remaining low-quality particles. Final reconstruction of the RBD-down trimeric spike was achieved through non-uniform refinement in C3 symmetry using a dataset of 142,853 particles. Model building was carried out manually in Coot, and real-space refinement was performed using the CCP-EM suite.

### X-ray structure determination of the BD55-1205 Fab bound to the SARS-CoV-2 BA.2.86 RBD

Codon-optimized cDNAs encoding the BD55-1205 Fab heavy chain (HC) with a C-terminal thrombin–His6–Strep tag (tHS) and the corresponding light chain (LC) were synthesized, subcloned into pcDNA3.4, and sequence-verified (GenScript). The SARS-CoV-2 BA.2.86 RBD (residues 330–528) was synthesized with the same C-terminal tHS tag, cloned into pcDNA3.4, and sequence-verified (GenScript). BD55-1205 Fab was produced by transient transfection of Expi293F cells (Thermo Fisher Scientific) using Turbo293 per the manufacturer’s instructions. Conditioned medium was harvested 5 days post-transfection and applied to Strep-Tactin® XT 4Flow® resin (IBA Lifesciences). Eluted material was exchanged into thrombin cleavage buffer and digested with thrombin, then passed over Ni–NTA resin to remove protease, cleaved tags, and uncleaved species. The Fab was polished by size-exclusion chromatography (SEC) on a Superdex 200 Increase 10/300 GL column (Cytiva) equilibrated in PBS.

BA.2.86 RBD was expressed in Expi293F GnTI− cells (Thermo Fisher Scientific) transiently transfected with Turbo293. Five days post-transfection, supernatants were purified on Strep-Tactin® XT 4Flow® resin, exchanged into thrombin cleavage buffer, and cleaved with thrombin. The reaction was cleaned up on Ni–NTA resin to remove protease, cleaved tags, and uncleaved RBD, followed by SEC on Superdex 200 Increase 10/300 GL in PBS.

Purified BD55-1205 Fab and BA.2.86 RBD were mixed at a 1:1.3 molar ratio (Fab:RBD), concentrated using a 10-kDa MWCO concentrator (Amicon), and injected onto a Superdex 200 Increase 10/300 GL equilibrated in 10 mM HEPES, pH 7.4, 150 mM NaCl to isolate the complex and exchange buffers. The BD55-1205 Fab–BA.2.86 RBD complex was concentrated with a 10-kDa MWCO device to 14.4 mg ml−1 for crystallization screening. Crystallization was carried out at 20 °C using an SPT Labtech Mosquito with commercial sparse-matrix screens (100 nl protein complex + 100 nl reservoir). Initial hits were obtained in Index HT (Hampton Research): 100 mM sodium acetate, pH 4.5, 25% (w/v) PEG 3350. Conditions were optimized to 100 mM sodium acetate, pH 4.5, 19% (w/v) PEG 3350 in hanging-drop plates with 0.5 μl protein complex and 0.75 μl reservoir. Crystals were cryoprotected in reservoir solution supplemented with 20% (v/v) glycerol, looped, and flash cooled.

Diffraction data were collected at the National Synchrotron Light Source II (Brookhaven National Laboratory) beamline 19-ID (NYX) using a Dectris Eiger2 9M XE detector. Data was processed using XDS^19^ and POINTLESS^20^ and AIMLESS^21^ to 2.5 Å. The crystal space group is P22121, with unit cell dimensions: a=54.9, b=111.5, c=300.8 (α=β=γ=90°). The structure was determined with molecular replacement using PDB entry 8XE9 with PHASER^22^ in CCP4 suite^23^. REFMAC5^24^ (CCP4 suite) was used for refinement and COOT^25^ for model building. Two BD55-1205 Fab–BA.2.86 RBD complexes are present in the asymmetric unit. Coordinates and structure factors have been deposited in the Protein Data Bank.

### In vitro expression of SARS-CoV-2 mRNA constructs

Expi293F cells (Thermo Fisher Scientific, Waltham, MA) were used for all *in vitro* transfection experiments. Cells were cultured in Expi293 Expression Medium (Thermo Fisher Scientific) and maintained in suspension at 37 °C, 8% CO_2_, with shaking at 125 rpm in a humidified incubator. Cells were seeded at 1 × 10^6^ cells/mL for all transfections and confirmed to be in log-phase growth with >97% viability at the time of use.

To assess in vitro expression, mRNA constructs encoding SARS-CoV-2 mRNA-1273 (JN.1 [control] and LP.8.1) and mRNA-1283 (mRNA1273.167 [front-run], JN.1 [control], LP.8.1 Codon Sequences 1–3) were transfected into Expi293F suspension cells (Thermo Fisher Scientific, Waltham, MA). Transfections were performed in 96-well deep-well plates using a 6-point, 2-fold serial dilution starting at 500 ng mRNA per 1 × 10^6^ cells, following the manufacturer’s protocol for the TransIT-mRNA transfection kit (Mirus Bio, Madison, WI). Cells were incubated at 37 °C, 8% CO_2_ with shaking at 1,000 rpm and harvested approximately 24 hours post-transfection.

Cell surface protein expression was evaluated by flow cytometry using binding to recombinant human ACE2 (hACE2) and mAb 1205 (RBD-specific), all of which contain human Fc regions. For mRNA-1283 constructs, additional binding was assessed for mAb 118 (NTD-specific). Samples were stained with an Alexa Fluor 647–conjugated goat anti-human IgG (Southern Biotech, Birmingham, AL) and Aqua viability dye (Thermo Fisher Scientific).

Flow cytometry was performed using an iQue3 (Sartorius) at a sip rate of 1.5 μL/s for 5 seconds per well. Data were collected with ForeCyt software (Sartorius) and analyzed by sequential gating on cells → singlets → viable cells → positive cells. Expression was quantified by calculating percent positive cells, median fluorescence intensity (MFI) of viable cells that is indictive of translation capacity, and Viable Cells MFI × percent positive for construct comparison. Graphs and statistical analyses were performed using GraphPad Prism (GraphPad Software).

### Sample Preparation and LC-MS/MS Acquisition

Samples were thawed and lysed with a combination of 5% SDS (Fisher Scientific, BP1311-1) in MS-grade water (Fisher Scientific, W6-4) containing protease inhibitors (Roche, 04 693 159 001) and sonication. Lysates were digested using a semi-automated digestion workstream using the Kingfisher Apex platform (Thermofisher Scientific) and MS-grade trypsin (Pierce, 90057). Following digestion, samples were acidified to a final concentration of 0.1% formic acid (Fisher Scientific, A117-50) before storage at 4C before analysis.

Acidified samples were separated using a 15cm × 75um Pepmap C18 column (Thermofisher Scientific, ES75150PN) coupled to a Vanquish Neo UHPLC system. Peptides were separated using the following gradient (A→B): 0-2% over 2 minutes, 2-27% over 21 minutes, 27-45% over 3 minutes, 45-95% over 2 minutes, followed by washing and equilibration. Data was acquired on an Exploris 480 Mass spectrometer (Thermofisher Scientific) using a data independent acquisition strategy. MS1 data was collected at 60K resolution followed by 32 sequential 8 m/z MS2 windows at 15k resolution. Normalized collision energy was set at 27% and MS2 AGC was set to 1000%. Data was analyzed using Spectronaut (Version 18.2) with a custom FASTA containing the Human Uniprot proteome (UP000005640) with the addition of the JN.1 and LP8.1 1273 amino acid sequence included.

### Animal studies

Animal studies were performed using eight-week-old, female BALB/mice (Charles River Laboratories). All immunizations were performed intramuscularly with either 0.5 or 1 µg of mRNA-LNP material or a PBS control in 50 µL of volume in a single hindleg. Serum was collected via submandibular bleed and terminal bleed was performed via cardiac puncture under isoflurane sedation. Following euthanasia, spleens were harvested for relevant studies. Eight cohorts of 32 animals each were used in these studies.

All experiments and procedures involving mice were conducted in accordance with Protocol 23-07-016, as approved by the Moderna’s Institutional Animal Care and Use Committee (IACUC). Activities performed adhere to the standards outlined by the NIH guidelines, Animal Welfare Act, and applicable US federal regulations. Euthanasia was performed using isoflurane inhalation followed by exsanguination under anesthesia and cervical dislocation, in accordance with the American Veterinary Medical Association (AVMA) Guidelines for the Euthanasia of Animals (2013 Report of the AVMA Panel on Euthanasia).

### Enzyme-linked immunosorbent assay

Assays were performed in 96-well microtiter plates (Thermo Fisher Scientific) coated with 100 μL of recombinant Wuhan-Hu-1 spike protein (S-2P). Plates were incubated at 4°C overnight and then blocked for 1 hour at 4°C using SuperBlock (Thermo Fisher Scientific). Sera were serially diluted in 5% BSA in TBS (TBS5; Boston Bioproducts, IBB-187), added to plates, incubated for 1 hour at 37°C, and then washed 4 times with TBS5. Goat anti-mouse IgG-HRP (Southern Biotech; Cat. #1030-05) was diluted in TBS5 then added and plates were incubated for 1 hour at 37°C. Plates were washed 4 times with TBS5 before the addition of TMB substrate (Thermo Fisher Scientific). Reactions were stopped by the addition of TMB stop solution (Sera Care; Cat. #5150-0021). Optical density measurements were taken at 450 nm, and titers were determined using a 4-parameter logistic curve fit in Prism Version 9 (GraphPad). Titers were defined as the reciprocal dilution at an OD of approximately 1 at 450 nm (normalized to a mouse standard on each plate).

### Vesicular stomatitis virus-based pseudovirus neutralization assay

Codon-optimized full-length spike genes (Wuhan-Hu-1 with D614G, BA.4/BA.5, XBB.1.5, JN.1, KP.2, KP.3.1.1, XEC, LP.8.1, LF.7.2.1, NB.1.8.1, and XFG) were cloned into a pCAGGS vector. To generate VSVΔG-based SARS-CoV-2 pseudovirus, BHK-21/WI-2 cells were transfected with the spike expression plasmid and infected by VSVΔG-firefly-luciferase as previously described (Whitt 2010 Journal of Virological Methods; PMID 20709108). A549 cells (ATCC; CCL-185) engineered to express human ACE2 and TMPRSS2 were used as target cells for the neutralization assay and maintained in DMEM supplemented with 10% fetal bovine serum. To perform the neutralization assay, serum samples were heat-inactivated for 45 min at 56ºC and serial dilutions were made in DMEM supplemented with 10% FBS. The diluted serum samples or culture medium (serving as virus only control) were mixed with VSVΔG-based SARS-CoV-2 pseudovirus and incubated at 37ºC for 45 min. The virus-serum mixture was subsequently used to infect A549-hACE2-TMPRSS2 cells for 18 h at 37ºC. Post infection, an equal volume of One-Glo reagent (Promega; E6120) was added to culture medium for readout using a PHERAstar FSX plate reader (BMG). The percentage of neutralization was calculated relative to the relative light units of the virus only control and subsequently analyzed using the four parameter logistic curve function in Prism 10 (GraphPad).

### T cell stimulation and intracellular cytokine staining

Following euthanasia, mouse spleens were collected and stored in RPMI10 media. Single-cell suspensions were prepared from BALB/c mouse spleens using a Fraunhofer tissue dissociator. After tissue dissociation, red blood cell lysis was performed for 2 minutes using ACK lysis buffer, followed by quenching with RPMI10 media and filtration through a 70 μm filter. Cells from each mouse were counted using a Cellaca MX high-throughput cell counter (Revvity Part. #MX-AOPI), resuspended in RPMI10 media.

After counting, splenocytes were centrifuged at 350×g for 5 minutes. Supernatant was removed and splenocytes were resuspended in 0.5 mL cold FBS at a concentration of 40-50×10^6^ cells/mL. Cold freezing media (0.5 mL; FBS+20% DMSO) was added drop by drop to the cell suspension. Cell suspension was then transferred to a prechilled cryovial and transferred to a Mr. Frosty freezing container. Splenocytes were frozen at −80ºC for 24 hours and then transferred to liquid nitrogen for long term storage.

For thawing, cryovials were placed in a Thawsome device (Medax, 1-9X-NIH1-S) in a 15-mL tube containing prewarmed RPMI10 media. Samples were centrifuged at 350×g for 5 minutes. Samples were washed with RPMI10 media and resuspended for cell counting. Cell counts were adjusted to 20×10×10^6^ live cells/mL. Splenocytes were incubated at 37°C with 5% CO_2_ for 6 hours in the presence of Protein Transport Inhibitor, 2 μg/mL of 15-amino-acid peptides overlapping by 11 amino acids encoding variants of interest (D614G, BA.4/BA.5, XBB.1.5, JN.1, KP.2, XEC, and LP.8.1), and CD107a PE (BD, Cat. #558661, clone 1D4B). Control conditions included DMSO (0.5% final, negative control) and PMA/Ionomycin (positive control).

After incubation, cells were washed with PBS and stained with Live/Dead Fixable UV Blue Dead Cell Stain (Invitrogen, Cat. #L34962) for 15 minutes at room temperature. Cells were then washed with FC stain buffer (PBS supplemented with 2% HI FBS and 0.05% sodium azide). Surface staining was performed with a cocktail of antibodies prepared in Brilliant Stain Buffer (BD, Cat. #566349), including FC Block (BD, Cat. #553142, clone 2.4G2), CD4+ BV480 (BD, Cat. #565634, clone RM4-5), CD4+ 4 BUV395 (BD, Cat. #740215, clone IM7), I-A/I-E AF488 (BD, Cat. # 562352, clone M5/114), and CD8+ BUV805 (BD, Cat. #612898, clone 53-6.7). The staining cocktail was added to the cells and incubated for 15 minutes at room temperature. Cells were then washed with FC stain buffer, fixed, and permeabilized using the BD Cytofix/Cytoperm kit (BD, Cat. #554714) according to the manufacturer’s instructions. After permeabilization, cells were washed with 1X Perm/Wash Buffer.

Intracellular staining was performed at 4°C for 20 minutes using a cocktail of antibodies in Brilliant Stain Buffer, including CD3 BV605 (BD, Cat. #564009, clone 17A2), IFN-γ BV421 (BD, Cat. #563376, clone XMG1.2), TNF-α BV711 (BD, Cat. #563944, clone MP6-XT22), IL-5 APC (BD, Cat. #554396, clone TRFK5), IL-4 PE-CF594 (BD, Cat. #562450, clone 11B11), IL-13 PE-Cy7 (Invitrogen, Cat. #25-7133-82, clone eBIO13A), and IL-2 BV785 (Biolegend, Cat. #503843, clone JES6-5H4). Finally, cells were washed with 1X Perm/Wash Buffer and FC stain buffer, filtered through a 30 μm filter in a 96-well plate, and resuspended in 0.5% PFA-FC stain buffer. Samples were acquired on an Aurora Spectral Flow Cytometer (Cytek Biosciences). T cell ICS analysis was subsequently performed using OMIQ. Lymphocytes were gated from the total population on an FSC-A vs SSC-A plot, followed by doublet exclusion using a singlet gate on an FSC-A vs FSC-H plot. Dead cells were excluded using a Live/Dead Blue vs SSC-A plot. CD3^+^MHCII− cells were gated from the live, single-cell population on an MHCII (IA-IE) vs CD3+ bivariate plot to exclude antigen-presenting cells. CD4^+^ and CD8^+^ T cell populations were separated on a CD8^+^ vs CD4^+^ bivariate plot. Cytokine expression (IFN-γ, TNF-α, IL-2, IL-4, IL-5, and/or IL-13) and the degranulation marker CD107a (CD8^+^ T cells only) were assessed within the CD4^+^ and CD8^+^ populations via co-expression with the T cell activation marker CD44 on a CD44 vs cytokine bivariate plot. Background cytokine expression from the no-peptide control condition (DMSO) was subtracted from that measured in peptide-stimulated samples for each individual mouse. Values ≤0.005 were thresholded to 0.0025, which represents a lower limit of detection that is 2-fold lower than the lowest observed positive signal.

## Acknowledgements

Cryo-EM data was collected at the Cryo-EM Facility at MIT.nano. We acknowledge beamline 19-ID (NYX) at NSLS-II, a U.S. DOE Office of Science User Facility at Brookhaven National Laboratory (Contract DE-SC0012704) and thank Kevin Battaile for X-ray data collection. We gratefully acknowledge all data contributors, i.e., the Authors and their Originating laboratories responsible for obtaining the specimens, and their Submitting laboratories for generating the genetic sequence and metadata and sharing via the GISAID Initiative, on which some of this research is based.

## Notes

### Competing Interest Statement

All authors are employees of Moderna and may own stock and/or stock options in the company. The authors declare no other competing interests.

