## Supplemental Figures for "mRNA-1273.251 and mRNA-1283.251 vaccines expressing SARS-CoV-2 variant LP.8.1 antigens broadly neutralize contemporary JN.1-lineage viruses"

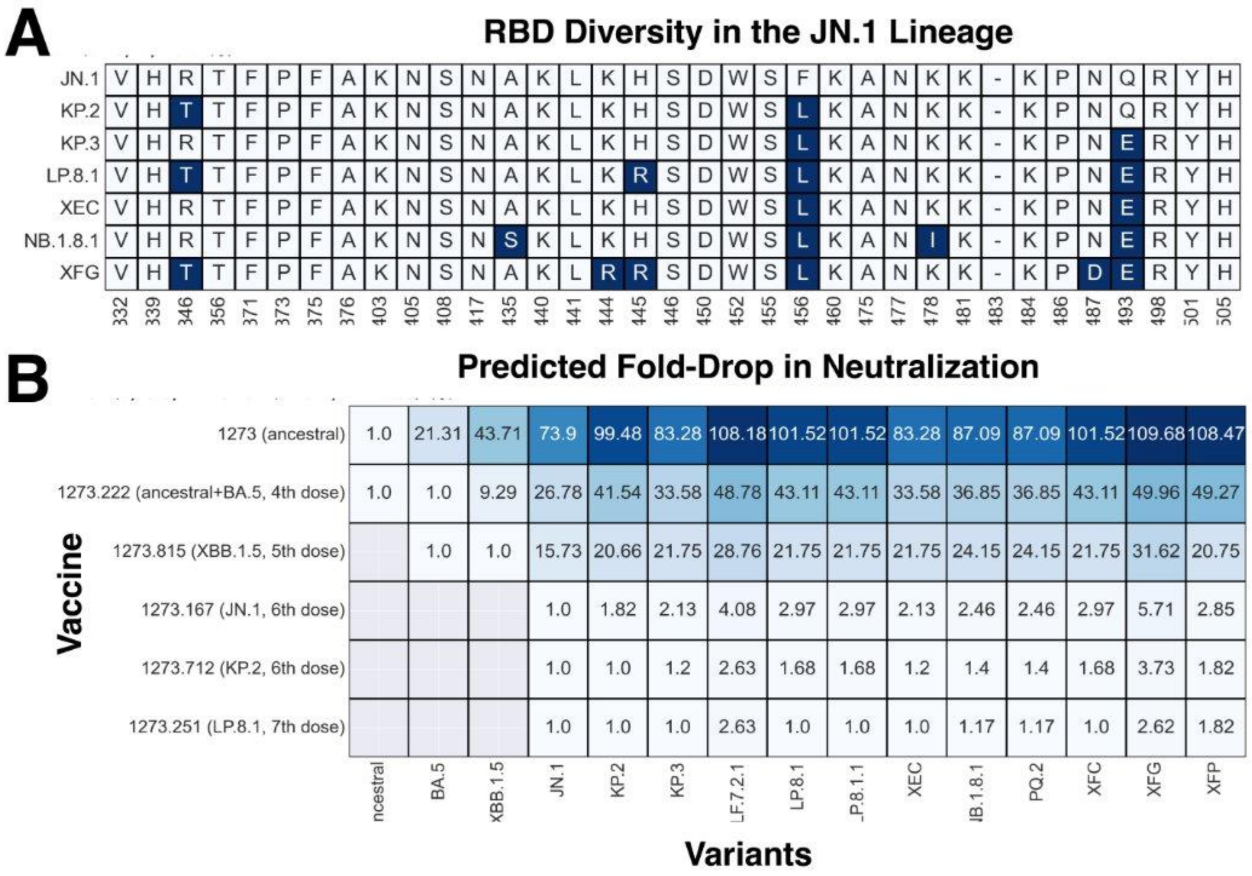


**Figure S1.** (A) S RBD mutations in JN.1 lineage variants. The x-axis indicates the amino acid position relative to the ancestral (Wuhan-Hu-1) strain. Mutations are indicated in different shades of blue. (B) The matrix lists the predicted immune escape for the same variants against different immunization backgrounds. The values inside the matrix are the predicted fold-drop in theID50 neutralizing antibody GMTs on the vaccine strain, derived from the statistical model described in Nasir et al.

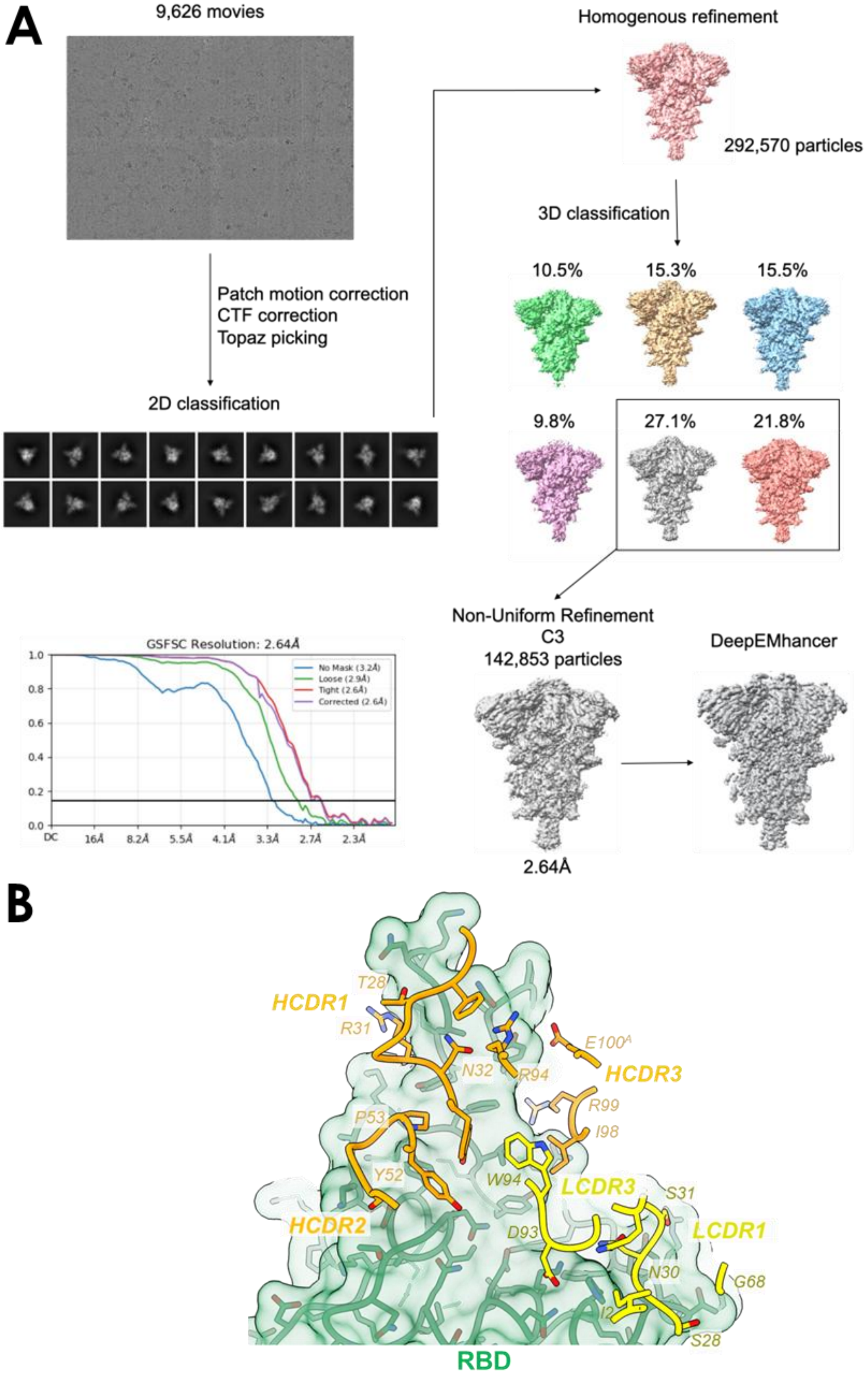
A
9,626 movies
Patch motion correction
CTF correction
Topaz picking
2D classification
Homogenous refinement
292,570 particles
3D classification
10.5%
15.3%
15.5%
9.8%
27.1%
21.8%
GSFSC Resolution: 2.64Å
No Mask (3.2Å)
Loose (2.9Å)
Tight (2.6Å)
Corrected (2.6Å)
1.0
0.8
0.6
0.4
0.2
0.0
DC
16Å
8.2Å
5.5Å
4.1Å
3.3Å
2.7Å
2.3Å
Non-Uniform Refinement
C3
142,853 particles
2.64Å
DeepEMhancer
B
HCDR1
T28
R31
N32
R94
E100A
HCDR3
R99
I98
P53
Y52
W94
LCDR3
S31
HCDR2
D93
LCDR1
N30
G68
S28
RBD

**Figure S2**. (A) Workflow description of Cryo electron microscopy refinement. Movies are captured of fields of particles and individual proteins are isolated using particle picking software. Particles are organized into 2D classifications based on visual properties and refined into a single 3D architecture. 3D classifications of the particles comprising that structure are broken out and refined to a final structure. Gold-standard Fourier Shell Correlation revealed the resolution of the structure to 2.64Å. (B) BA.2.86 RBD / BA55-1205 Fab interface. Heavy-chain (*HCDR1–HCDR3*) and light-chain (*LCDR2–LCDR3*) CDR loops are depicted above the RBD surface. Antibody 1205 residues contacting the RBD—defined as any antibody atoms within 4.0 Å of an RBD atoms—are shown.

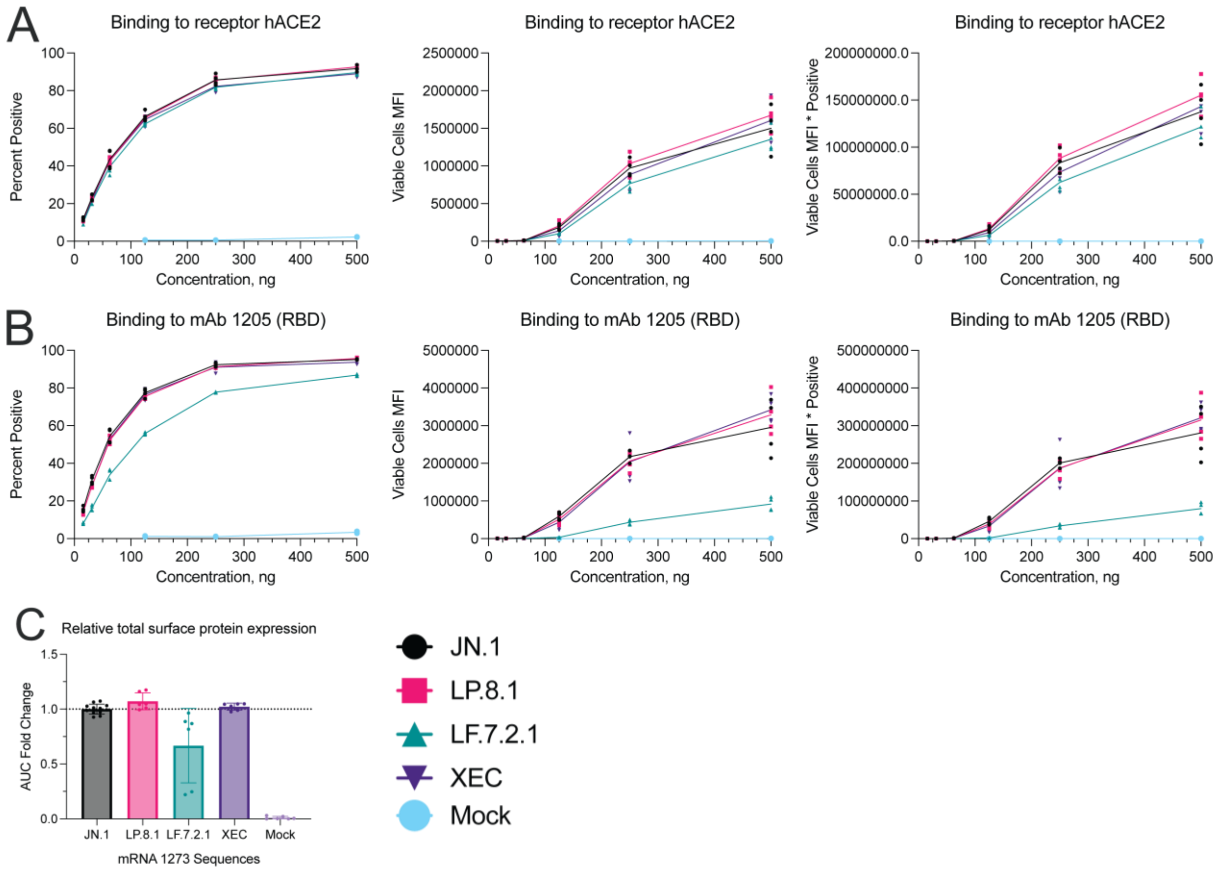


**Figure S3**. *In vitro* expression curves of Expi293F cells transfected with mRNA expressing mRNA-1273 designs of SARS-CoV-2 antigens. Curves represent multiple masses of mRNA transfected and graphs describe the percentage of positively expressing cells, median fluorescence intensity (MFI) of viable cells and the product of these two readouts from left to right after detection with the (A) hACE2 receptor or (B) an RBD-targeting monoclonal antibody. (C) Cells were then processed and examined via LC-MS for relative expression in a reagent agnostic manner.

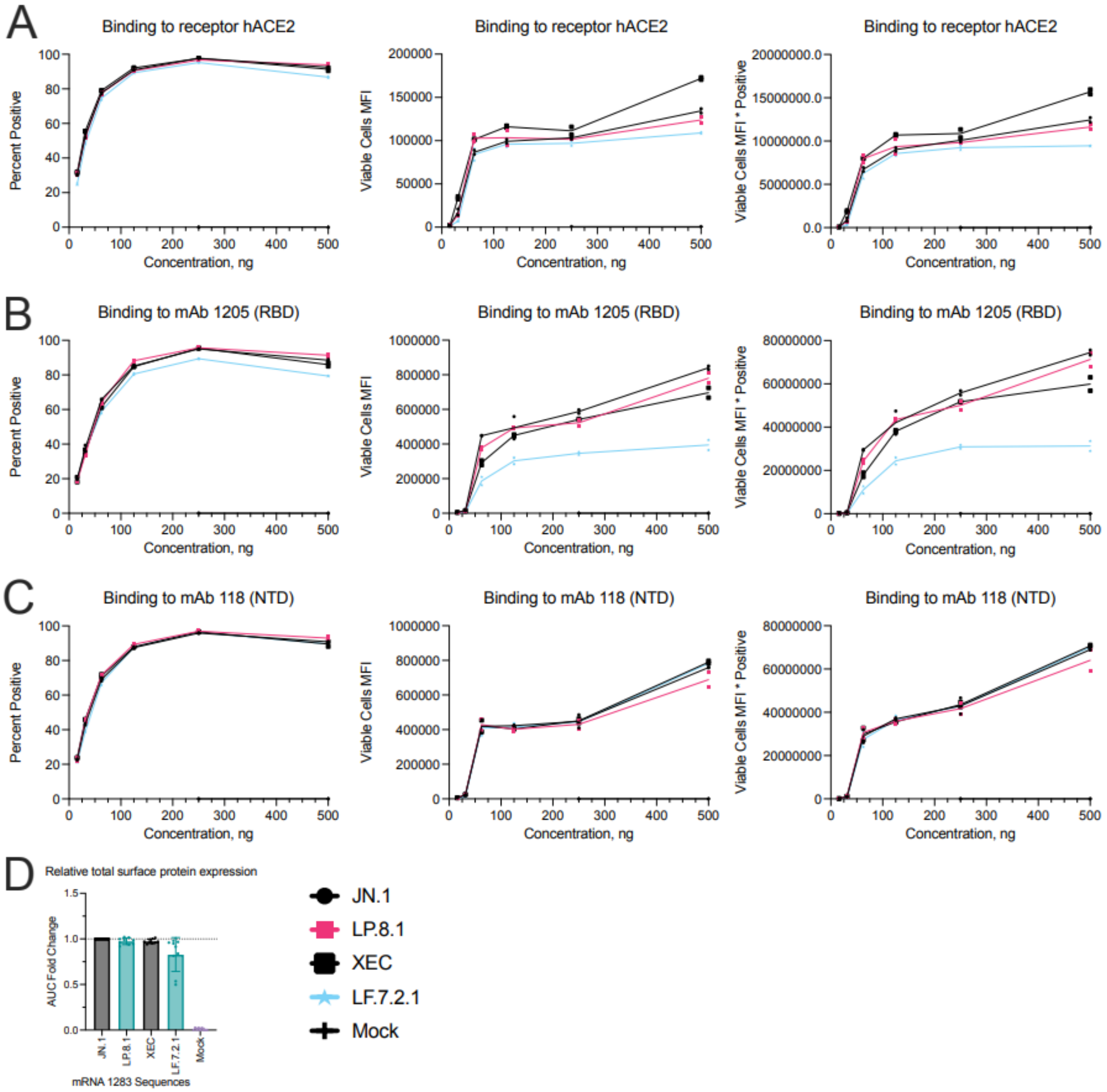


**Figure S4**. *In vitro* expression curves of Expi293F cells transfected with mRNA expressing mRNA-1283 designs of SARS-CoV-2 antigens. Curves represent multiple masses of mRNA transfected and graphs describe the percentage of positively expressing cells, median fluorescence intensity (MFI) of viable cells and the product of these two readouts from left to right after detection with the (A) hACE2 receptor, (B) an RBD-targeting monoclonal antibody, or (C) an NTD-targeting monoclonal antibody. (D) Cells were then processed and examined via LC-MS for relative expression in a reagent agnostic manner.

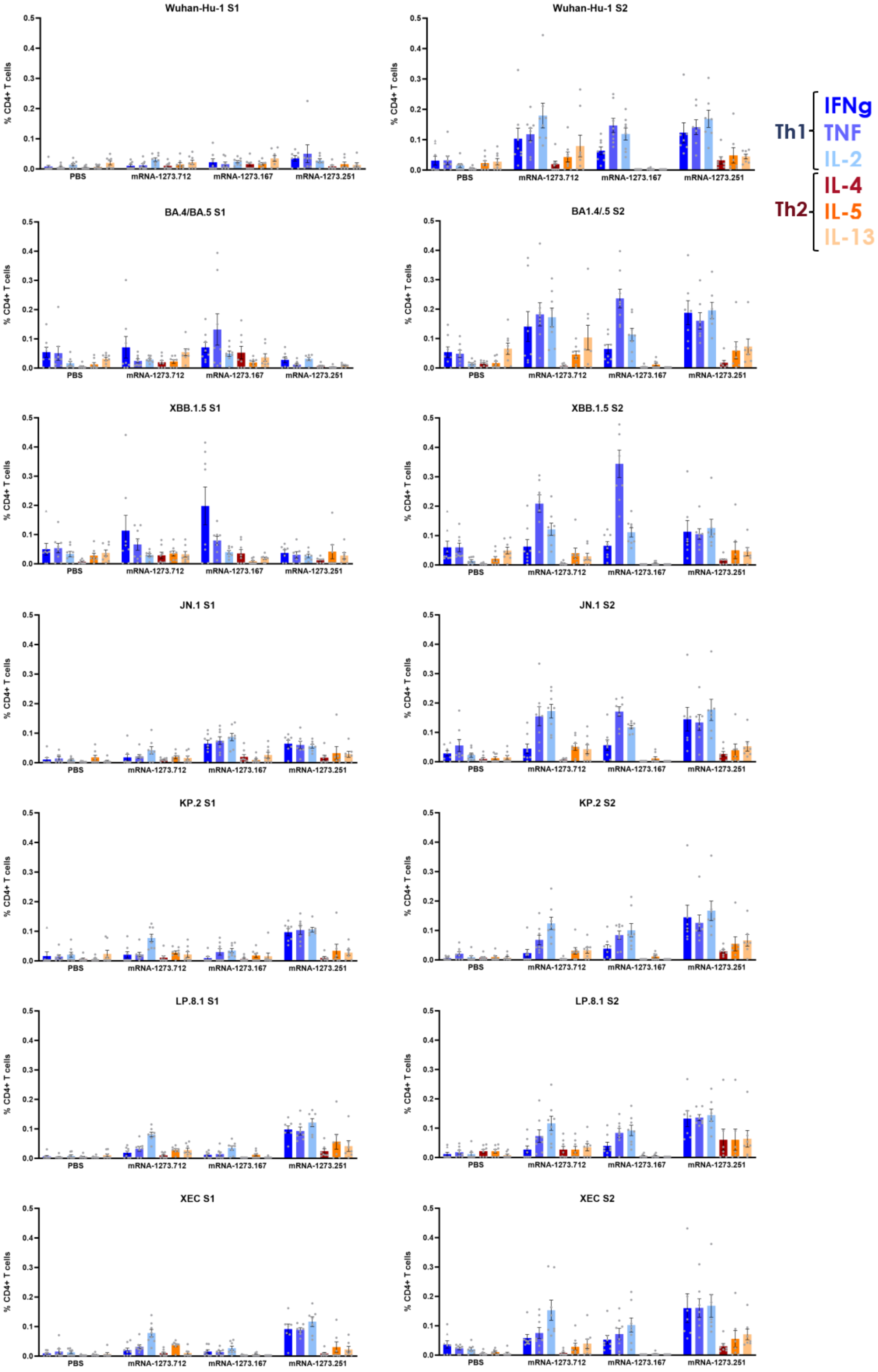
Wuhan-Hu-1 S1
Wuhan-Hu-1 S2
BA.4/BA.5 S1
BA1.4/.5 S2
XBB.1.5 S1
XBB.1.5 S2
JN.1 S1
JN.1 S2
KP.2 S1
KP.2 S2
LP.8.1 S1
LP.8.1 S2
XEC S1
XEC S2
% CD4+ T cells
PBS
mRNA-1273.712
mRNA-1273.167
mRNA-1273.251
Th1
IFNg
TNF
IL-2
Th2
IL-4
IL-5
IL-13

**Figure S5.** Full Th1 and Th2 cytokine panels are shown for CD4 T-cells reactive to peptide pools representing the S1 (left) or S2 (right) domains of the SARS-CoV-2 spike protein for each variant tested. Th1 cytokines are represented by shades of blue and Th2 cytokines by shades of red.

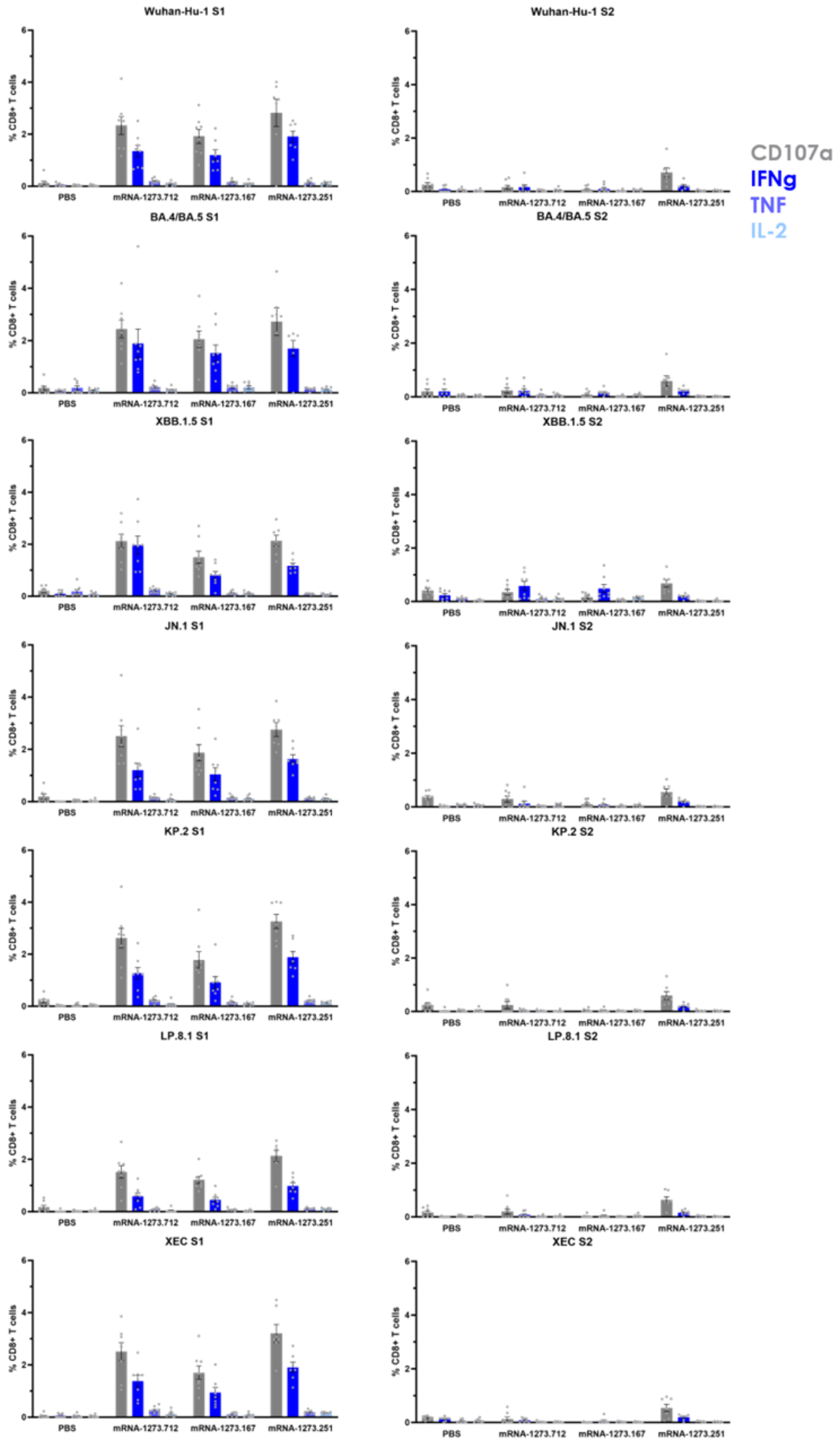

Wuhan-Hu-1 S1
Wuhan-Hu-1 S2
BA.4/BA.5 S1
BA.4/BA.5 S2
XBB.1.5 S1
XBB.1.5 S2
JN.1 S1
JN.1 S2
KP.2 S1
KP.2 S2
LP.8.1 S1
LP.8.1 S2
XEC S1
XEC S2
% CD8+ T cells
PBS
mRNA-1273.712
mRNA-1273.167
mRNA-1273.251
CD107a
IFNg
TNF
IL-2

**Figure S6.** Full cytokine panels are shown for CD8 T-cells reactive to peptide pools representing the S1 (left) or S2 (right) domains of the SARS-CoV-2 spike protein for each variant tested.
